# The Xist locus on the X chromosome of males regulates cognition and metabolism despite minimal expression

**DOI:** 10.64898/2026.11.11.766005

**Authors:** Nilla Sivakumar, Dan Wang, Dena B. Dubal, Samira Abdulai-Saiku

## Abstract

The long non-coding RNA Xist is the master regulator of X chromosome inactivation (XCI) in females, but its functional significance in males, where Xist expression is normally minimal, is unknown. Here we investigated whether the Xist locus influences male physiology and brain function across the lifespan using mice carrying a targeted Xist deletion (ΔXist). Xist deletion produced a small further reduction in the very low, residual Xist signal in aged male hippocampus. Metabolically, Xist deletion lowered fasting blood glucose in old, not young, males, without affecting body weight. In the brain, Xist deletion modestly improved spatial memory at 24 hours in the Morris water maze in young males, with a similar trend at middle age in an interactive place avoidance task, independent of anxiety-like behavior or locomotor activity. This early cognitive benefit did not persist with aging and motor function on the rotarod was unaffected by genotype at any age. Together, these findings indicate that the Xist locus exerts age- and domain-specific effects on male metabolic and cognitive functions despite near-absent baseline Xist expression. These data identify an unexpected role for a canonically female-associated regulatory locus in males and raise the possibility that Xist-associated mechanisms contribute to male brain function and metabolism.

## Introduction

X chromosome inactivation (XCI) transcriptionally silences one X chromosome in female mammalian cells to achieve dosage compensation between XX and XY individuals^1^. XCI is initiated and maintained by Xist, a long non-coding RNA (lncRNA) that coats the inactive X and recruits silencing machinery in females^2,3^. Beyond dosage compensation and sex determination, emerging evidence implicates X chromosome biology in aging and disease across tissues^4^.

In males, which carry a single active X chromosome, Xist is normally expressed at very low levels and its functional significance remains largely unexplored. However, the Xist locus can exert regulatory effects independent of robust transcript expression, potentially through chromatin-level mechanisms at the locus itself^5^. Moreover, Xist expression can also become activated in male cells in disease, including male cancers^6^ and in autoimmune conditions like lupus^7^. These observations raise the possibility that Xist or the Xist locus itself has biological functions in males despite minimal transcript expression.

Aging provides an additional context in which Xist regulation may be relevant. Recent work in our lab and others demonstrates that aging activates escape from silencing and increases Xist expression in the female mouse brain^8,9^, various other mouse tissues^10^, and several human tissues^11^. Whether Xist also contributes to brain or systemic phenotypes across the male lifespan is unknown. Here, we address this question by characterizing the metabolic and cognitive consequences of Xist deletion in male mice across the lifespan, from young adulthood through old age.

### Characterization of Xist deletion in young and old male mice

Since the Xist locus may have regulatory effects^3,12^, Xist expression can emerge in males^6,7^, and aging increases Xist in the female mouse brain^8^, we tested whether the Xist locus affects male aging and cognition. We generated males with Xist deletion (ΔXist; Fig. 1a) on their X chromosome.

**Figure 1:**
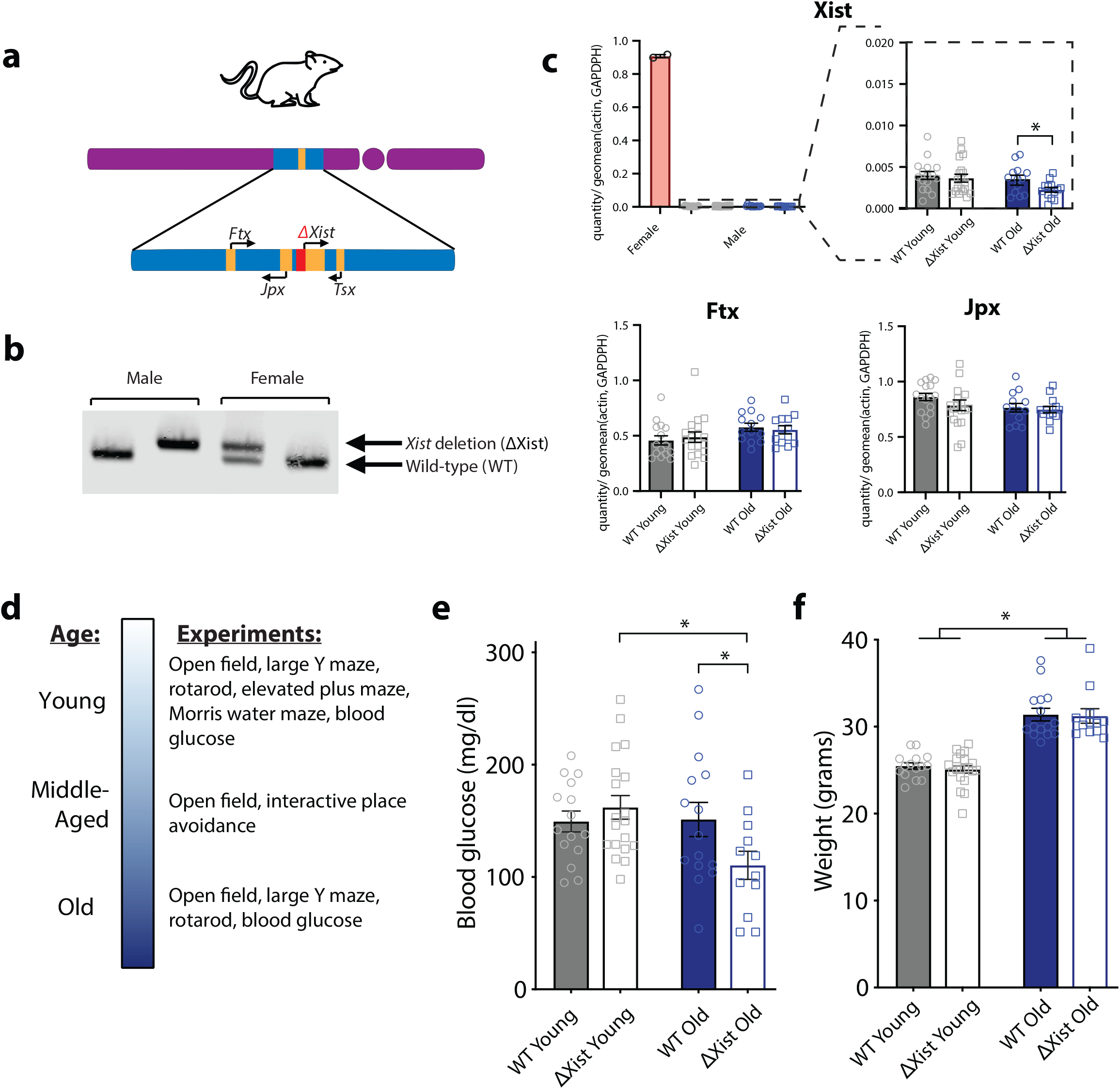
Characterization of Xist deletion in young and old male mice. **a**, Male mice underwent Xist knockout (ΔXist). Diagram of the mouse adapted from IMG Visual Icons/Shutterstock. **b**, Genotyping confirmation of Xist deletion in males and females. **c**, Quantification of mRNA levels for key X-inactivation factors *Xist, Ftx*, and *Jpx* in young and old male hippocampus. *Xist* gene expression was nearly undetectable in males (upper right) compared to females (upper left) and was further reduced in old ΔXist compared to old WT mice (^*^p = 0.0318, Welch’s t-test). (Young, age 4-8 months: WT n = 16, ΔXist n = 19; Old, age 22-24 months: WT n = 14, ΔXist n = 13). **d**, Timeline of experiments conducted over the lifespan. **e**, Xist deletion decreased fasted blood glucose levels in old animals but not in young ones (^*^p = 0.0486, Welch’s t-test.) (Young, age 4-8 months: WT n = 15, ΔXist n = 19; Old, age 22-24 months: WT n = 15, ΔXist n = 12). **f**, Aging, but not Xist deletion, increased body weight (age: ^****^p < 0.0001, two-way ANOVA). (Young, age 4-8 months: WT n = 15, ΔXist n = 19; Old, age 22-24 months: WT n = 15, ΔXist n = 12).

Following confirmation of Xist deletion in males and females by genotype (Fig. 1b), we quantified mRNA levels for Xist and other X-inactivation factors Ftx and Jpx in young and old male hippocampus. Ftx and Jpx did not differ between groups. As expected^2^, Xist gene expression was nearly undetectable in males compared to females.

Nonetheless, Xist deletion decreased the low residual expression in old male mice (Fig. 1c). In contrast to findings in females^8,9^, Xist did not increase in brain aging in males (Fig. 1c).

We next assessed the metabolic and behavioral consequences of Xist deletion across age in males (Fig. 1d). Xist deletion decreased fasting blood glucose levels in old, but not in young male mice (Fig. 1e), indicating an age-dependent metabolic effect. As anticipated^13^, old mice weighed more than young mice; however, Xist deletion did not alter body weight at either age (Fig. 1f).

### Xist deletion modestly improves memory in young male mice and trends toward improved memory in middle-aged male mice

We next asked whether deleting Xist in males influences brain function. We first used the elevated plus maze assay to assess anxiety-like behavior and activity (Fig. 2a). Time spent in the open arm, an anxiety-like measure, did not differ between groups (Fig. 2b). Distance traveled also did not differ (Fig. 2c), indicating that Xist deletion did not alter anxiety-like behavior or general activity.

**Figure 2:**
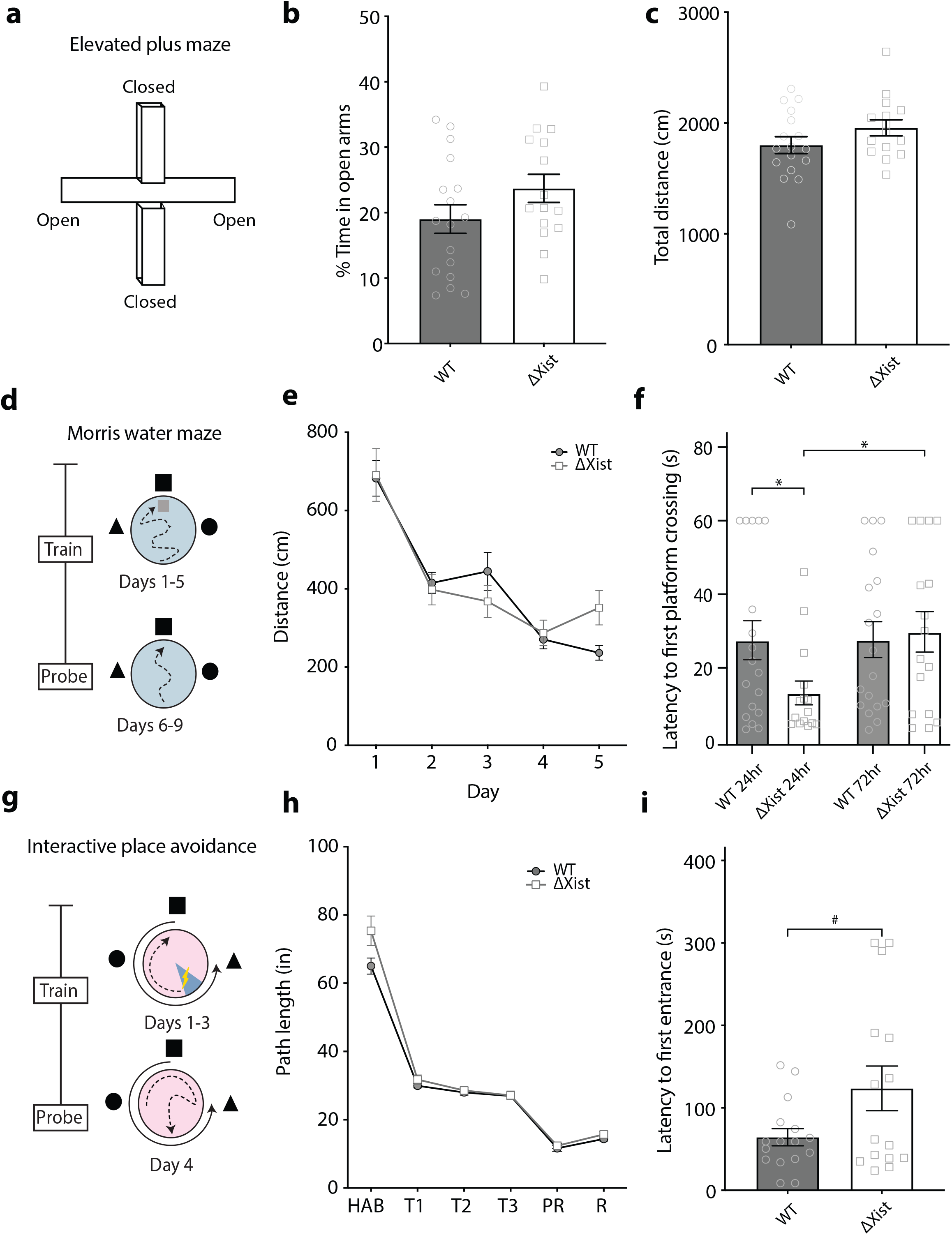
Xist deletion modestly improves memory in young male mice and trends toward improved memory in middle-aged male mice. **a**, Schematic of the elevated plus maze assay. **b**, Time spent in the open arm, an anxiety-like measure, did not differ between the groups (Age 4-8 months; WT: *n* = 17, ΔXist: *n* = 15). **c**, Total distance traveled in the elevated plus maze did not differ between groups (Age 4-8 months; WT: *n* = 17, ΔXist: *n* = 15). **d**, Schematic of the Morris water maze paradigm. **e**, Spatial learning in the hidden trials, measured by distance traveled to find the platform, did not differ between the groups (age by genotype: *p* = 0.2578, two-way ANOVA) (Age 4-8 months; WT: *n* = 18, ΔXist: *n* = 16). **f**, Xist deletion improved memory of the platform location in the 24h memory probe (^*^*p* = 0.0290, Welch’s t-test), but not 72 hours (*p* = 0.7762, Welch’s t-test). ΔXist mice performed better at 24h than 72h (^*^*p* = 0.0177, paired t-test) whereas WT mice showed no difference between the two probes (Age 4-8 months; WT: *n* = 18, ΔXist: *n* = 16). **g**, Schematic of interactive place avoidance (IPA) experiment. **h**, Distance traveled during IPA did not differ between the groups (*p* = 0.8263, two-way ANOVA) (Age 10-12 months; WT: *n* = 18, ΔXist: *n* = 15). **i**, Xist deletion tended to increase latency to enter the former shock zone, indicating a trend toward better memory (#*p* = 0.0565, Welch’s t-test) (Age 10-12 months; WT: *n* = 16, ΔXist: *n* = 15).

We then used the Morris water maze to assess spatial learning and memory (Fig. 2d). Spatial learning in the hidden trials, measured by distance traveled to find the platform, did not differ between the groups (Fig. 2e). In contrast, Xist deletion improved memory of the platform location in the 24h memory probe; this early effect did not persist to 72 hours (Fig. 2f). Thus, Xist deletion enhanced early memory retention in males.

To determine whether Xist deletion improves cognition in middle-age mice, we used the interactive place avoidance (IPA) task (Fig. 2g). Distance traveled during IPA did not differ between the groups (Fig. 2h). Xist deletion tended to increase latency to enter the former shock zone, indicating a trend toward better memory (Fig. 2i). Together, these findings indicate that Xist deletion confers a modest improvement to memory in young male animals, with a similar direction of effect in middle-age, independent of anxiety-like behavior or locomotor activity.

### Aging impairs memory and motor function in male mice, with memory decline more evident with Xist deletion

Given the modest cognitive benefit of Xist deletion earlier in the life of males, we next asked whether this effect persists into old age or alters age-related cognitive decline. We first used the open field assay to assess context-dependent spatial learning and memory across age (Fig. 3a). Spatial habituation and dishabituation did not differ by genotype across the lifespan (Fig. 3b), indicating Xist deletion did not broadly alter this form of context-dependent behavior.

**Figure 3:**
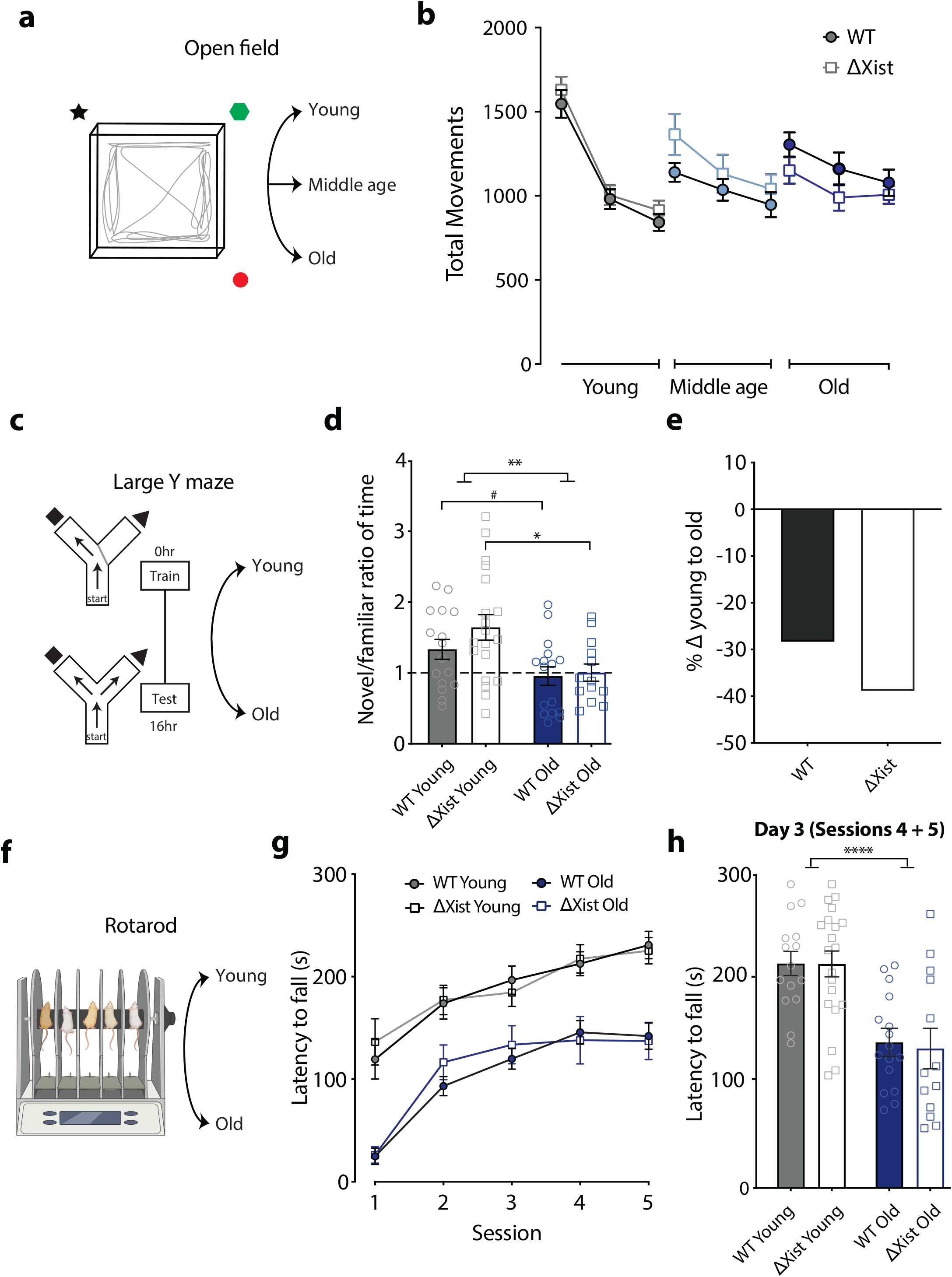
Aging impairs memory and motor function in male mice, with memory decline more evident with Xist deletion. **a**, Schematic of the open field assay to assess context-dependent spatial learning and memory. **b**, Spatial habituation and dishabituation did not differ across lifespan by genotype. (genotype by age by day: *p* = 0.9206, three-way ANOVA) (Young, age 4-8 months: WT n = 18, ΔXist n = 15; Middle-aged, age 10-12 months: WT n = 18, ΔXist n = 15; Old, age 22-24 months: WT n = 17, ΔXist n = 13) **c**, Diagram of large Y maze to evaluate spatial learning and memory in young (age 4– 8 months) and old (age 20–24 months) male mice. **d**, Aging decreased memory (Age: ^**^*p* = 0.0053, two-way ANOVA), with a significant decrease in ΔXist mice (^**^*p* = 0.0063, Welch’s t-test) and a trend toward decrease in WT mice (#*p* = 0.0582, Welch’s t-test). (Young, age 4-8 months: WT *n* = 16, deltaXist *n*= 19; Old, age 22-24 months: WT *n* = 16, deltaXist *n* = 13) **e**, Percentage of change in memory from the young to old life stage in both experimental groups in the Y-maze. **f**, Schematic of rotarod to evaluate motor coordination and learning in young (age 4– 8 months) and old (age 20–24 months) male mice. Image adapted from Biorender. **g**, Aging, but not Xist deletion, increased latency to fall from rotarod at all points during rotarod training (age: ^****^p < 0.0001, session by age by genotype: p = 0.3477, three-way ANOVA) (Young, age 4-8 months: WT n = 15, ΔXist n = 20; Old, age 22-24 months: WT n = 16, ΔXist n = 13). **h**, Aging, but not Xist deletion, increased latency to fall from rotarod on final day of assay. (age: ^****^p < 0.0001, age by genotype: p = 0.8359, two-way ANOVA). (Young, age 4-8 months: WT n = 15, ΔXist n = 20; Old, age 22-24 months: WT n = 16, Δxist n = 13).

We then used a large Y maze paradigm to evaluate spatial learning and memory in young (4-8 months) and old (20-24 months) male mice (Fig. 3c). Aging decreased memory, with a significant decrease in ΔXist mice and a trend toward decrease in WT mice (Fig. 3d). We further calculated the percentage change in memory from the young to old age in each genotype (Fig. 3e) and observed a greater percentage change in memory in ΔXist compared to WT mice. Thus, despite modest memory benefit observed at younger ages, Xist deletion did not preserve memory in old males and tended toward a more pronounced age-related decline in this measure.

Finally, we used a rotarod assay to determine whether Xist deletion altered age-related changes to motor function (Fig. 3f). Aging decreased latency to fall across rotarod training, whereas Xist deletion had no effect (Fig. 3g). Similarly, aging, but not Xist deletion, decreased latency to fall on the final day of testing (Fig. 3h). Taken together, these findings indicate that effects of Xist deletion are age- and domain-dependent. Xist deletion in males modestly enhances memory earlier in life but does not protect against age-related cognitive or motor changes in old male mice.

## Discussion

Here, we demonstrate Xist deletion from the X chromosome in male mice affects metabolism and cognition in an age-dependent manner. Xist deletion decreased fasting blood glucose in old males, improved short-term spatial memory in young males, and tended to decrease memory in old age. Xist deletion did not alter motor function.

Together, these findings suggest that the Xist locus, even when largely transcriptionally silent in males, influences biological phenotypes in health and aging. These findings raise the question of how the Xist locus could influence metabolism and cognition when Xist is hardly transcribed in males? Several precedents point to transcript-independent mechanisms. LncRNA loci can regulate neighboring genes through promoter, enhancer-like, transcriptional, or splicing features rather than just by the mature RNA^14,15^. For example, the Ftx locus influences Xist expression independently of its RNA products^16,17^ in females. Xist sits within a regulatory landscape organized by CTCF-dependent chromatin architecture^18-21^, so its deletion could rewire local chromatin folding in cis^22^. Alternatively, rare cells may express Xist, as in male cancers^6^ and lupus^7^, where Xist ribonucleoprotein complexes can reprogram immune cells in males^23^. Finally, global Xist deletion may have developmental consequences in adulthood. Resolving these possibilities will require cell type-specific and tissue-specific analyses across development and aging.

More broadly, these findings raise the possibility that regulation of the X chromosome has an underappreciated role in male health. Deleting Xist improved cognition and metabolism even though Xist is largely silent in males, suggesting that regulatory mechanisms within the X itself could cause biological effects without Xist expression. Males would be particularly sensitive to such effects because the X is enriched for genes affecting cognition^24^ - and with only a single X, they lack a second allele to buffer disruptions in X-linked regulation, as illustrated by X-linked disorders^25^.

Improved memory in young males with Xist deletion suggests that Xist or Xist-associated mechanisms constrain aspects of male brain function. The loss of this benefit in aging suggests that Xist regulation and its consequences may shift across the lifespan, consistent with age-induced changes in X-inactivation stability and Xist expression in females^8,9,11^. Xist activation in male cancers^6^ and lupus^7^, altered X-inactivation in T cells in sex-biased autoimmunity^26^, and even mosaic loss of the Y chromosome ^27,28^ further illustrate the dynamic regulation of sex chromosomes across health, aging and disease^4^. Together, our findings identify an unexpected role for the Xist locus in males and raise the possibility that Xist and related regulatory mechanisms contribute to male vulnerability and resilience across the lifespan.

## Materials and Methods

### Animals

All animal studies were conducted under protocols approved by the Institutional Animal Care and Use Committee at the University of California, San Francisco (UCSF). Animals were maintained on a 12-hour light/dark cycle under standard environmental conditions (temperature: 19–23 °C; relative humidity: 30–70%) with unrestricted access to food and water. Mice were group-housed in cohorts of five per cage, with the exception of animals undergoing Morris water maze testing, which were individually housed during the course of testing. All behavioral assessments were conducted during the light phase.

To investigate the effects of Xist deletion, we generated mice carrying a global Xist deletion and compared them to non-transgenic wildtype littermate controls. Briefly, 129-Xist^tm2Jae^/Mmnc mice obtained from the Mutant Mouse Resource and Research Centers were crossed with Zp3-Cre mice. The resulting F2 offspring were subsequently backcrossed onto a C57BL/6J background, with congenicity confirmed by genetic testing. Male mice were evaluated across a battery of behavioral and cognitive assessments across the lifespan. All testing apparatus and arenas were thoroughly cleaned with 70% ethanol between subjects, with the exception of the water maze. Experimenters remained blinded to genotype and group assignments throughout all experiments.

### Morris Water Maze

The water maze was run according to previously published protocols^29^. We used a circular pool (122 cm diameter) filled with white, opaque water kept at 21 ± 1 °C, with a 14 cm^2^ escape platform hidden 2 cm below the water surface. Before hidden platform training began, each mouse completed two pretraining trials in which it swam down a straight channel to reach a visible platform. During hidden platform training, the platform stayed in one place while the starting position varied from trial to trial. Mice were given four trials per day for eight consecutive days. In probe trials, the platform was taken out and mice swam freely for 60 seconds, and latency to reach the platform’s former location was measured. After the probe trials, mice were tested on a visible platform version of the task, in which the platform was flagged with a 15 cm vertical cue.

### Open Field

Open-field testing followed previously published procedures^29^. Mice were habituated to the testing room for one hour and then allowed to explore for 10 minutes. The arena was a clear plastic enclosure (41 × 30 cm), and locomotor activity was measured by beam breaks using an automated Flex-Field/Open Field Photobeam Activity System (San Diego Instruments). Visual cues marked three of the chamber walls. For repeated testing across the lifespan, each animal was placed in the same chamber it had used previously.

### Elevated Plus Maze (EPM)

The EPM was performed as described previously^29^, with dim lighting during both habituation and testing. Mice were habituated to the room for one hour beforehand. Each mouse was placed at the center of the maze facing an open arm and allowed to explore for 10 minutes. The Kinder Scientific Elevated Plus Maze and MotorMonitor system recorded the distance traveled and the proportion of time spent in open versus closed arms.

### Two-Trial Large Y Maze

The two-trial large Y maze, which has visual cues at the end of each arm, was run as described previously^29^. During training, mice could access only the familiar arm while the novel arm was blocked. Sixteen hours later, the mice were returned to the maze with all three arms open and explored freely for five minutes. AnyMaze software was used to track the time spent in the novel versus familiar arms, and a novel-to-familiar exploration ratio was then calculated.

### Interactive Place Avoidance (IPA)

Mice were habituated to the testing room for one hour before testing in a square white enclosure (40 × 40 cm) under dim lighting. In each session, animals were placed on a rotating disk containing a predefined electrified zone, with cues on the surrounding walls, and could move about freely. Video recordings were analyzed with the CleverSys TopScan Automated Behavior Analysis System (v.3.0) to extract the times at which mice entered the electrified zone.

### RT–qPCR

Hippocampi were dissected from animals perfused with PBS. They were then flash-frozen. RNA was extracted using the Qiagen RNEasy Mini Kit (74104) and cDNA was prepared using Applied Biosystems’ TaqMan Reverse Transcription Reagents (N8080234). Primers were designed via the NCBI Primer-BLAST tool and synthesized by Integrated DNA Technologies.

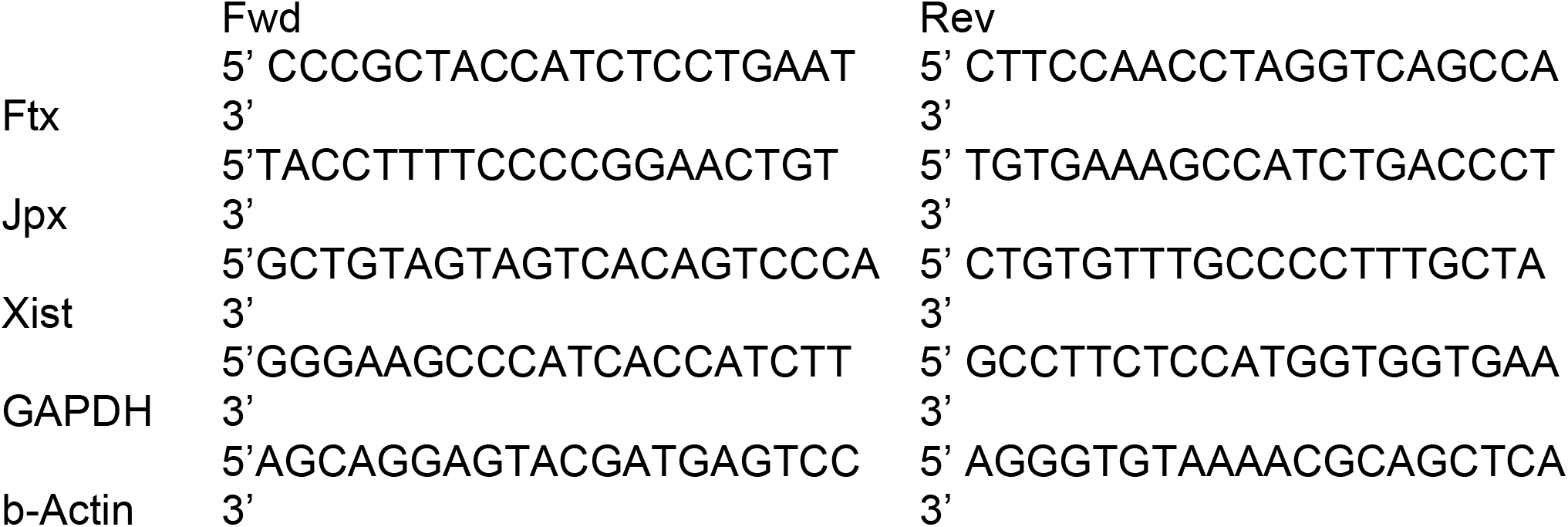

### Statistical Analysis

All experimenters were blinded to genotype throughout data collection and analysis. Statistical analyses were performed using GraphPad Prism (v.11.0) for t-tests and two-way ANOVAs, and RStudio (v. 2026.01.2) for mixed-model ANOVAs and post-hoc comparisons. All tests were two-tailed unless otherwise specified. Comparisons between two unpaired group means were made using Welch’s unpaired t-tests, and differences among multiple means were evaluated using two-way ANOVA, unless otherwise noted. Post-hoc comparisons were corrected for multiple comparisons using the Bonferroni–Holm method in R, controlling the family-wise error rate at α = 0.05 (rounded to two decimal places), unless otherwise indicated. Morris water maze and open-field data were analyzed using a mixed-model ANOVA that accounted for repeated measures. Outlier exclusion criteria were defined a priori as values falling more than 2 standard deviations above or below the group mean, ensuring unbiased removal of extreme observations. Error bars throughout depict the standard error of the mean (SEM). Statistical significance was set at *p* ≤ 0.05 (rounded to two decimal places) and marked by asterisk (^*^), and statistical trends were set at *p* ≤ 0.10 and marked by pound symbols (#).

## Acknowledgements

We thank Barbara Panning for Xist deletion discussions and guidance in generating Xist knockout mice, S. Kalantry for Zp3-Cre mice used to generate the Xist deletion animals, C. Chen for mouse colony maintenance, and the Gladstone Behavioral Core for assistance with behavioral studies. Primary funding for the study was by the National Science Foundation (to N.S.), NIH grants RF1AG079176-01 and RF1AG068325 (D.B.D.), American Federation for Aging Research (D.B.D.), the Bakar Aging Research Institute (D.B.D.), the Simons Foundation (1018027 to S.A.-S. and 811225SPI to D.B.D.), and philanthropy (D.B.D.).

## Declaration of Interests

D.B.D. is an inventor on issued US patents and applications involving klotho, X chromosome factors, PF4 and cognitive improvement. S.A.S. is an inventor on a patent application involving X chromosome factors and cognitive improvement. D.B.D. is a co-founder of Jocasta Neurosciences and chairs the scientific advisory board.

### Declaration of Generative AI and AI-Assisted Technologies in the Manuscript Preparation Process

In preparation of this Perspective, the authors used ChatGPT (OpenAI) to assist with language editing, clarity, and organization of the manuscript. All scientific content, interpretations, and conclusions were created, reviewed and verified by the authors, who take full responsibility for the accuracy and integrity of the final manuscript.

## Notes

### Competing Interest Statement

The authors have declared no competing interest.

